## Supplementary figures with legends for "Scheduled feeding improves behavioral outcomes and reduces inflammation in a mouse model of Fragile X syndrome"

S. Fig. 1

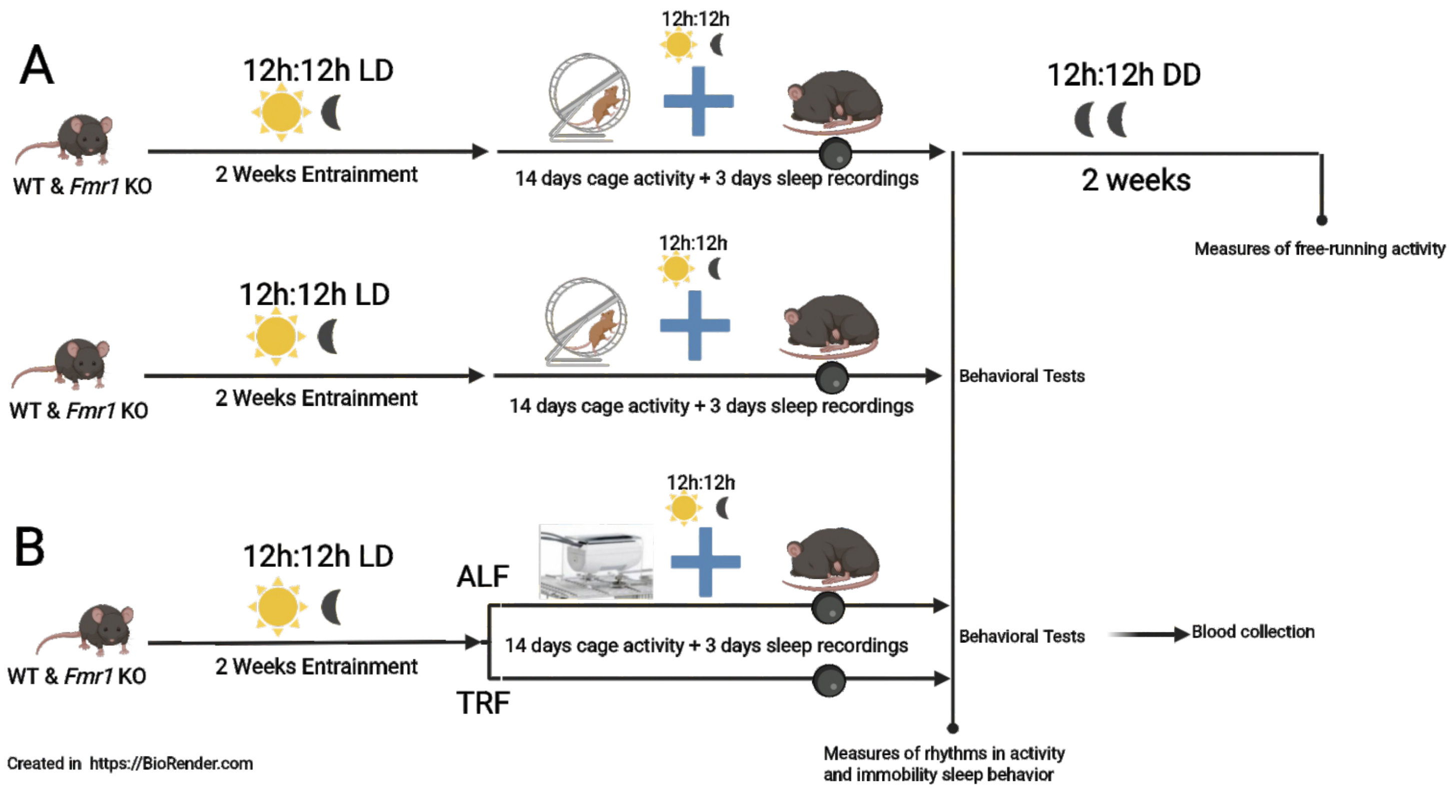


**Supplementary Figure 1:** Graphic presentation of the Experimental design. (**A)** Mice were entrained to a 12h:12h Light-Dark (LD) cycle followed by 14 days of cage activity and 3 days of immobility sleep recordings, then the animals were released in constant darkness (DD) to measure their activity rhythms driven by the endogenous clock (free-running activity) and not by external cues. Behavioral tests were conducted after the sleep/wake cycles recordings in a different group of mice. (**B**) A different cohort of WT and *Fmr1* KO mice after entrainment to the LD cycle for 2 weeks was divided in two groups, one was held on *Ad Libitum* feeding (ALF) and one on a scheduled feeding regimen (time-restricted feeding, TRF). Mice were allowed to freely eat for 6 hours between ZT15 and ZT 21. After cage activity and immobility sleep were recorded, the mice were tested for social and repetitive behaviors. Blood was collected in the early light phase.

### S. Fig. 2

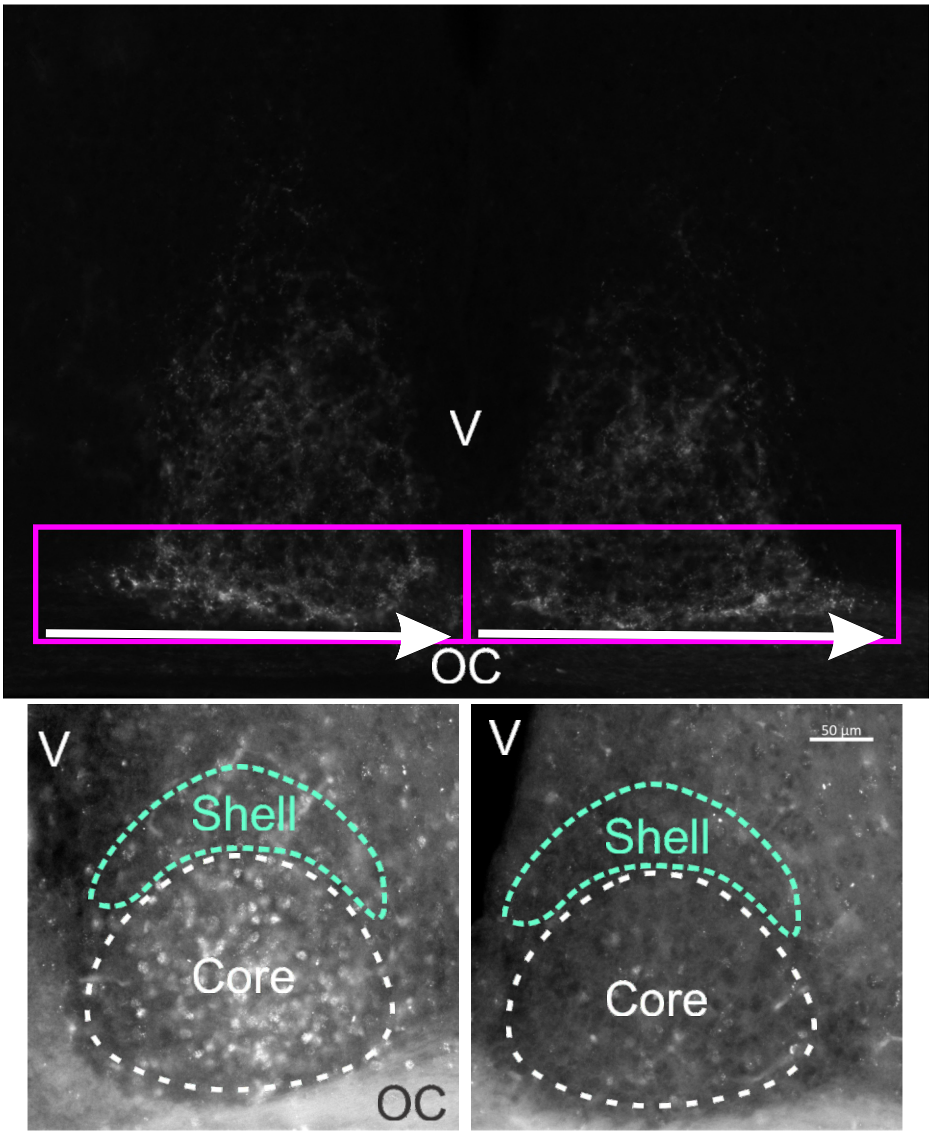


**Supplementary Figure 2: (A)** The distribution of the Cholera Toxin fluorescent signal was obtained for each left and right SCN using the Profile Plot Analysis feature of ImageJ. A rectangular box of fixed size (415.38µm x 110.94 µm, width x height) was created to include the entire ventral part of the SCN. A column plot profile was generated whereby the x-axis represents the horizontal distance through the SCN (lateral to medial for the left SCN and medial to lateral for the right SCN, as indicated by the arrows) and the y-axis represents the average pixel intensity per vertical line within the rectangular box. **(B**) Images with outlined the shell (green) and core (white) of the SCN, the master circadian clock, located in the anterior hypothalamus: OC= Optic Chiasm, V = third ventricle

### S. Fig. 3

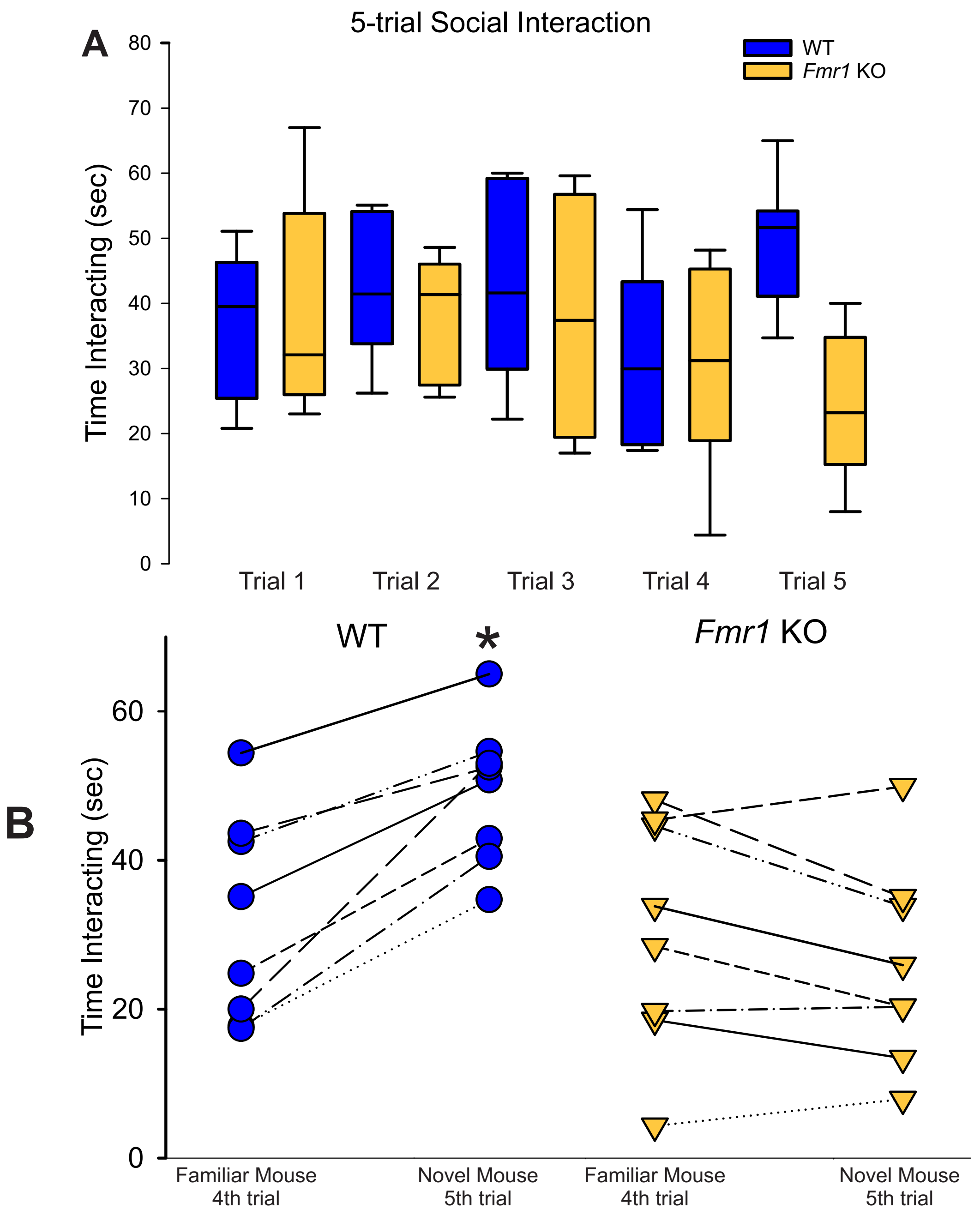


**Supplementary Figure 3:** To assess social recognition memory, Fmr1 KO and WT mice underwent a five-trial social interaction paradigm in a neutral open-field arena. Each trial lasted 2 minutes and was separated by a 5-minute inter-trial interval. During trials 1 through 4, the test mouse was exposed to the same unfamiliar conspecific (mouse A) enclosed within a wire cup to permit olfactory but limited tactile interaction. In trial 5, a novel conspecific (mouse B, stranger) was introduced. Time spent investigating the stimulus mouse (defined as sniffing or directing the nose toward the enclosure in close proximity) was manually scored. A progressive decrease in investigation time across trials 1–4 reflects habituation, while a significant increase in trial 5 indicates dishabituation and suggest intact social recognition memory. (**A**) Box plots: time spent by the testing mice interacting in each of the 5 trials. The boundary of the box closest to zero indicates the 25th percentile, the line within the box marks the median and the boundary of the box farthest from zero indicates the 75th percentile. The whiskers above and below the box indicate the 90th and 10th percentiles. (**B**) Plots showing the interaction time for each mouse during $4^{th}$ (familiar mouse) and $5^{th}$ (novel mouse) trial. The WT mice exhibited significant increases in the time spent interacting with the stimulus mouse in the $5^{th}$ trial as compared to the between the $4^{th}$ (paired *t*-test $T_{(14)} = 2.604$, P =0.021), while no differences were observed for the KO mice between the $4^{th}$ and the $5^{th}$ trial ($T_{(14)} = 0.624$, P = 0.542).

### S. Fig. 4

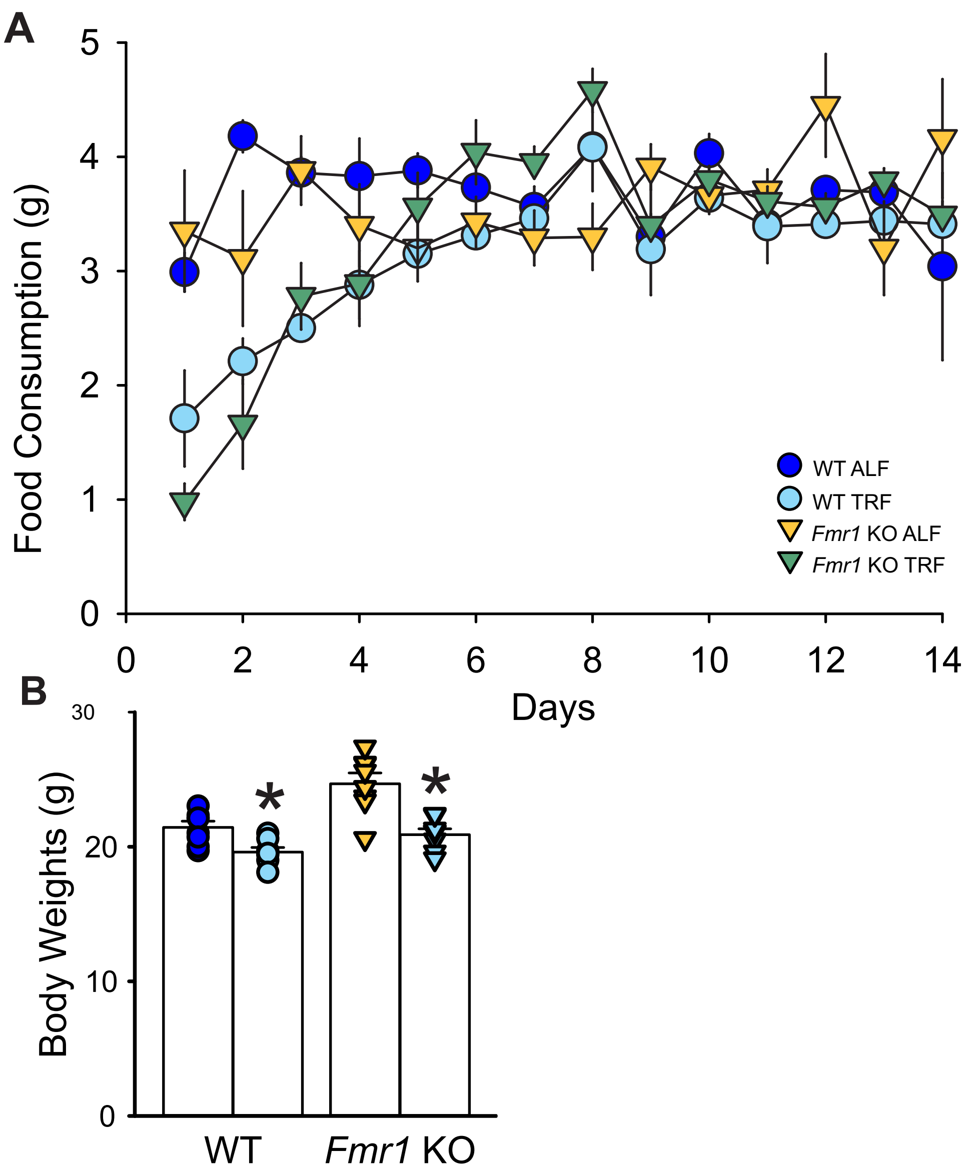


**Supplementary Figure 4: (A)** Food intake was similar across genotypes and conditions. Over the two weeks of scheduled feeding, no significant differences were found in the total amount of food consumed between genotypes ($F_{(1, 31)}$ = 3.086, P = 0.090) or feeding schedules ($F_{(1, 31)}$ = 0.307, *P* = 0.584). If we just analyzed the food consumed over the first three days, there were no effects of genotype ($F_{(1, 31)}$ = 2.737, P = 0.109), albeit the TRF groups consumed less ($F_{(1, 31)}$ = 85.912, *P* < 0.001). **(B)** Prior to TRF, the mice had similar weights (WT: 22.1 ± 0.4 g; KO: 23.1± 0.4 g; $t_{(30)}$ = -1.748, *P* = 0.090). After two weeks on TRF, both WT and Fmr1 KO mice exhibited lower weights than their counterparts on ALF (F = 30.551; *P* < 0.001). Two-Way Anova followed by the Holm-Sidak's multiple comparisons test. (**P* < 0.01)
